# Important anatomical, age-related, and species considerations regarding ocular fibulin-3 (EFEMP1) analysis

**DOI:** 10.1101/2022.09.28.509587

**Authors:** Steffi Daniel, John D. Hulleman

## Abstract

**Purpose:** Fibulin-3 (F3) or EFEMP1 is a secreted extracellular matrix glycoprotein implicated in several ocular diseases. Little is known about the native biology of this protein. Thus, our study aims to determine expression and localization characteristics of F3 utilizing a range of mammalian species as well as F3-associated changes with age.

**Methods:** Gene expression analyses for fibulins as well as immunohistochemistry for F3 were conducted in ocular tissue from mice, pigs, non-human primates (NHPs), and humans (n = 3-5). Age-based F3 expression study along with changes in ECM remodeling enzymes was also evaluated in mice.

**Results:** Within the mouse retina, F3 staining was consistent throughout the entirety of the retina (far-periphery, mid-periphery, and central), being enriched in the ganglion cell layer and inner nuclear layer (INL). However, in humans, the F3 staining pattern was quite unique; enriched in the RPE, INL, and outer nuclear layer (ONL) in the peripheral retina, but then shifting to predominantly outer plexiform layer (OPL) staining in the central retina and macula with waning RPE immunoreactivity approaching the fovea. We demonstrate that F3 expression in the mouse retina significantly increases with age, and the levels of extracellular F3 degrading enzymes produced by the RPE and retina (e.g., Mmp2 and Htra1) decrease with age.

**Conclusions:** These findings demonstrate that F3 has distinct species-dependent as well as ocular region-specific expression and localization patterns. We also show that F3 and ECM enzyme dynamics favor F3 accumulation in the retina and RPE with increasing age.

## INTRODUCTION

Fibulin-3 (F3) or epidermal growth factor (EGF)-containing fibulin-like extracellular matrix protein 1 (EFEMP1) is a secreted extracellular matrix (ECM) glycoprotein, belonging to the fibulin family of proteins^1^. F3, like many of the other members of its protein family, is widely expressed throughout the human body and plays an important role in regulating the structural and physiological integrity of basement membranes and the ECM^2–4^. Accordingly, loss of F3 expression (or loss of F3 function) is associated with reduced pelvic organ support, inguinal hernias, advanced bone age, ECM disruptions, and development of a Marfan-like syndrome^5–10^.

Conversely, and somewhat paradoxically, increases in F3 copy number/expression have been associated with a higher chance of development of ocular diseases such as age-related macular degeneration (AMD)^11^, ^12^, the leading cause of blindness in the elderly in industrialized nations^13^. Moreover, select autosomal dominant mutations in F3 appear to cause a range of diverse and devastating vision disorders including juvenile or primary open angle glaucoma^14–16^, generalized retinal degeneration^17^, and the rare AMD-like disease, Doyne honeycomb retinal dystrophy/Malattia Leventinese (DHRD/ML)^18^.

Given the wide range of human diseases in which F3 is involved, it is important for researchers to utilize appropriate model systems that accurately recapitulate F3-related human physiology and pathophysiology. For example, from a systemic disease perspective, F3 knockout mice mimic many of the same phenotypes^6^ observed in humans who lack F3 expression or function^7, 8^. Moreover, mice expressing the Arg345Trp (R345W) mutation that causes DHRD/ML primarily form age-dependent sub-retinal pigmented epithelium (RPE) basal laminar deposits (BLamDs)^19–21^ that are reminiscent of early AMD disruptions^22, 23^. Yet, these deposits do not resemble the extensive drusen typically observed in DHRD/ML patients^24^. The reasons for this discrepancy are not clear – whether such differences are due to inherent anatomical dissimilarities, such as a thinner Bruch’s membrane and elastin layer in mice (likely facilitating better transport and diffusion of macromolecules between the RPE and choroid vasculature)^25^, or whether lower levels of F3 expression in the mouse RPE^26^ could be to blame. Surprisingly, there are virtually no studies that compare and contrast F3-related ocular expression and protein localization information amongst mice, the most commonly used laboratory model to study visual disorders, and higher order mammals such as non-human primates (NHPs), or even humans themselves.

To bridge this gap in knowledge, in this study, we have thoroughly and rigorously evaluated similarities and differences in F3 transcription and localization across multiple ocular tissues implicated in F3-related disease (i.e., the trabecular meshwork ring, neural retina, and RPE/choroid) isolated from mice, pigs, NHPs, and humans. In complementary studies, we paralleled these findings with F3 immunofluorescence using a knockout-validated F3 antibody, and determined how F3 production/accumulation changes with advanced age and in diseases such as AMD. Ultimately, our results demonstrate the surprising and dynamic nature of F3 expression/localization which is dependent on the retinal area visualized, species used, and age. Integration of these factors, along with subject sex^27^, is necessary when interpreting F3-related observations in the eye, especially those related to pathophysiology of F3-associated ocular diseases, such as retinal degeneration and possibly glaucoma.

## MATERIALS AND METHODS

### Mice

C57BL/6J female mice were obtained from Jackson Laboratory (Bar Harbor, ME) at 18, 6, and 2 months of age. Additional C57BL/6 wildtype (17, 6 and 2 months) as well as F3 wildtype and knockout mice^26^ (15 months, males and females) from separate breeding protocols were also obtained. To ensure consistency, mice were analyzed within each of their respective groups. All mice were maintained in 12 h/12 h light/dark cycle and supplied with food and water *ad libitum*. All experiments were conducted in accordance with the Association for Research in Vision and Ophthalmology (ARVO) Statement of the Use of Animals in Ophthalmic and Vision Research and the UT Southwestern Medical Center (UTSW) Institutional Animal Care and Use Committee (IACUC) guidelines.

### Non-human primate (NHP) eyes (Olive baboon)

Normal NHP eyes (male and females between 1 and 12 years of age) (Table S1) were obtained from the Southwest National Primate Research Center (San Antonio, TX). Maximum death to preservation time was 4 h. One eye from each pair was preserved in RNALater solution at 4°C and used for quantitative PCR (qPCR), while the other was fixed in 10% formalin and used for histology.

### Human donor eyes

Normal adult donor eyes (between 65 and 85 years of age) (Table S2) with no known history of retinal diseases or corneal transplants were obtained from Lion’s Eye Institute (Tampa Bay, FL). Maximum death to preservation time was 6 h. One eye from each pair was preserved in RNALater solution (Invitrogen #AM7020; Carlsbad, CA, USA) and used for quantitative PCR, while the other fixed in Davidson’s solution (2 parts formalin, 3 parts ethanol, 1 part glacial acetic acid, and 3 parts water) and used for immunofluorescence. Three OCT frozen slides each from two early AMD donors were obtained from the Curcio Lab (University of Alabama at Birmingham, Table S3).

### Porcine eyes (Hampshire, Duroc, and Red Wattle hogs)

Adult porcine eyes ages (males and females between 6 months to 1 year of age) were obtained from Huse’s Country Meats Inc. (Malone, TX). Maximum death to preservation time was 4h. One eye from each pair was preserved in RNALater solution at 4°C and used for qPCR, while the other was fixed in 10% formalin and used for histology.

### qPCR analysis

RNA isolation for all tissues were carried out using the Aurum Total RNA isolation kit (BioRad #732-6820; Hercules, CA, USA). Samples were reverse-transcribed to cDNA using qScript cDNA Supermix (Quantabio #101414-106; Beverly, MA, USA). Available TaqMan probes (Applied Biosystems; Waltham, MA, USA) were used for qPCR reactions as follows: Mice: fibulin-1 (*Fbln1*, Mm00515700_m1), fibulin-2 (*Fbln2*, Mm00484266_m1), fibulin-3 (*Efemp1*, Mm00524588_m1), fibulin-4 (*Efemp2*, Mm00445429_m1), fibulin-5 (*Fbln5*, Mm00488601_m1), Mmp2 (Mm00439498_m1), Timp3 (Mm00441826_m1), Htra1 (Mm00479887_m1). Humans and NHPs: fibulin-1 (*FBLN1*, Hs00972609_m1), fibulin-2 (*FBLN2*, Hs00157482_m1), fibulin-3 (*EFEMP1*, Hs00244575_m1), fibulin-5 (*FBLN5*, Hs01056636_m1). Porcine: fibulin-3 (*EFEMP1*, Ss04326723_m1), fibulin-4 (*EFEMP2*, Ss06901209_m1), fibulin-5 (*FBLN5*, Ss04805024_m11) and samples were run on a QuantStudio 6 Real-Time PCR system (Applied Biosystems #4485694) in technical and biological replicates (n = 3-5). Relative abundance was calculated by comparing expression to β-actin [mice (*Bact*, Mm02619580_g1), humans and NHPs (*BACT*, Hs01060665_g1), and pigs (*BACT*, Ss03376563_uH) in each respective tissue.

### H&E histology

All eyes were processed into paraffin-embedded sections (Histo Pathology Core, UT Southwestern). H&E staining was performed, and images of the sections were taken at 10x and 20x magnification using Leica DMI3000 B manual inverted fluorescence microscope (Leica Microsystems, Buffalo Grove, IL).

### Immunohistochemistry

All fixed eyes were processed into paraffin-embedded sections, deparaffinized, and subjected to antigen retrieval (Histo Pathology Core, UT Southwestern). Slides were washed 2 × 5 min in 1x Tris-buffered saline (TBS) followed by pretreatment in 0.025% Triton X-100/1xTBS with gentle agitation (2 × 5 min). Sections were blocked in blocking solution consisting of 10% normal goat serum (New Zealand origin, #16210064, Gibco, Waltham, MA) with 1% bovine serum albumin (Fisher Bioreagents, Waltham, MA, BP1600-100) in TBS for 2 hours at room temperature (RT) in a humidified chamber. Slides were drained and incubated in primary antibodies diluted in blocking solution overnight at 4°C. The antibodies used were as follows: anti-fibulin-3 antibody (1:200, Santa Cruz, #sc-365224, Dallas, TX), anti-fibulin-3 antibody (1:100, Chemicon Millipore, #MAB1763, St. Louis, MO), anti-fibulin-3 antibody (1:100, GeneTex, #GTX111657, Irvine, CA), anti-fibulin-3 antibody (1:100, ProSci, #5213, Poway, CA,), fibulin-3 blocking peptide (5:1 to anti-fibulin-3 antibody, Prosci # 5213P). The next day, slides were rinsed with 0.025% Triton X-100/1xTBS 2 × 5 min and incubated in goat anti-rabbit AlexaFlour488 secondary antibody (1:1000, Life Technologies, #A27034, Carlsbad, CA) at RT for at least 2 h followed by 4’,6-diamidino-2-phenylindol (DAPI) staining at RT for 20 min. Sections were then rinsed with 1xTBS 3 × 5 min and mounted with Vectashield (Vector Laboratories, Newark, CA). Images were taken at 25x or 63x magnification using a Leica TCS SP8 confocal microscope.

## RESULTS

### Mice and higher order mammals have distinct patterns of fibulin gene expression in the neural retina and RPE

In previous studies, we were surprised to find that F3 expression levels in the neural retina (NR) of C57BL/6 mice were consistently higher than those found in the RPE/choroid^26^, the primary site of F3-driven retinal pathology^18–20^. We reconfirmed these important qPCR data (Fig. 1A) and expanded the same analyses to pigs, NHPs, and humans (Fig. 1B-D), utilizing all the commercially-available TaqMan fibulin probes for each species. Amongst the species analyzed, mice were the only animal where F3 expression was significantly higher (an average of 2-fold) in the NR compared to the RPE/choroid (Fig. 1A-D, *** p < 0.001), in keeping with our past observations. Fold increases in RPE/choroid F3 expression vs. NR expression ranged from 3-fold in pigs, to 2-fold in NHPs, to 10-fold in humans (Fig. 1B-D), all of which were significant (* = p < 0.05, ** = p < 0.01). Interestingly, in tissues with high levels of F3 expression (e.g., RPE/choroid from pig, NHPs, and humans), little expression of other fibulin family members was observed, indicating that F3 is the primary fibulin component in these tissues (Fig. 1B-D). In contrast, in tissues with low F3 expression (e.g., RPE/choroid from mice), high levels of other fibulins (fibulin-1, fibulin-4, and fibulin-5) were found (Fig. 1A), potentially as a compensatory mechanism. Nonetheless, these F3 expression differences already prognosticate likely F3 protein localization variation amongst the tested species.

**Figure 1.**
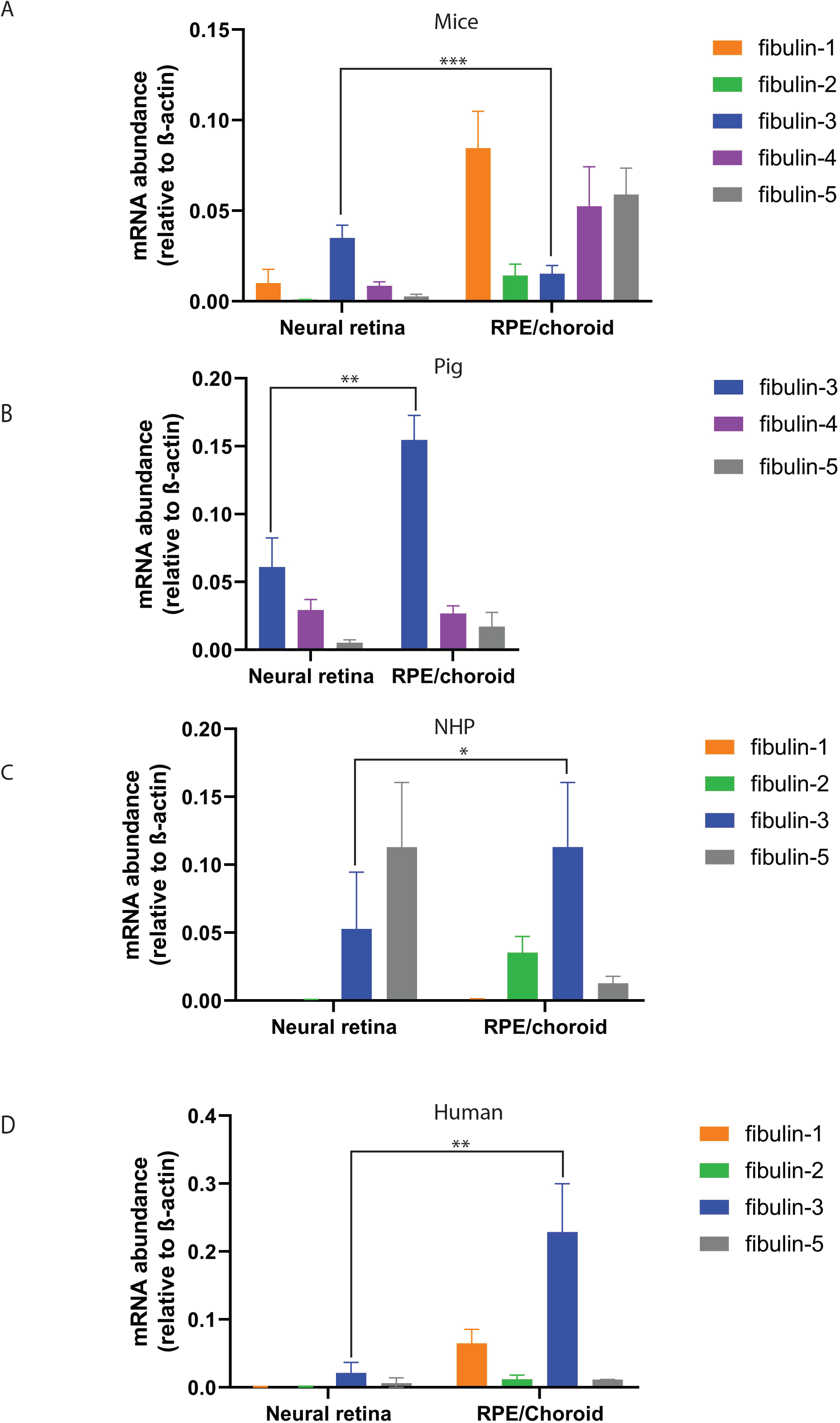
Gene expression profile of fibulins across species. (**A**) Expression levels of fibulin 1–5 in the NR and RPE of mice relative to β-actin (n = 3) (F3, p≤0.001). (**B**) Expression levels of fibulin 3–5 in the NR and RPE of pigs relative to β-actin (n =4) (F3, p≤0.01). (**C**) Expression levels of fibulin 1–3 and 5 in the NR and RPE of NHPs relative to β-actin (n = 4) (F3, p≤0.05). (**D**) Expression levels of fibulin 1–3 and 5 in the NR and RPE of humans relative to β-actin (n = 5) (F3, p≤0.01). Unpaired t test. Mean ± SD.

### Ocular distribution and localization of F3 protein varies between species

During our endeavor to better define F3 protein localization in the eye, we found that many available F3 antibodies produced a significant amount of background staining, even in fibulin-3 knockout (KO) mice (Fig. S1F-H). Moreover, in instances where we did not have a F3 KO control (i.e., in pigs, NHPs, and humans), it was difficult to gauge the specificity of staining across species, without the use of a blocking peptide. Thus, throughout our studies, we used an antibody from ProSci (cat# 5213) that showed no staining in F3 KO mice (Fig. S1E) and was also available with a corresponding blocking peptide (cat# 5113P). Consistent with our observations at the transcriptional level, within mice, F3 is found primarily in the ganglion cells layer (GCL) and inner nuclear layer (INL) with no clear signal originating from the RPE layer (Fig. 2A, Fig. S2A, B), consistent with previous difficulties detecting wild-type F3 in the RPE, even in aged mice^28^. Porcine ocular cross-sections revealed a unique, retina-wide, almost equal distribution of F3, including in the GCL, inner plexiform layer (IPL), INL, outer plexiform layer (OPL), outer nuclear layer (ONL), photoreceptors (PR), as well as RPE (Fig. 2B, Fig S2C, D). In contrast, distribution of F3 in the NHP and human ocular cross-sections exhibited patterns of localization particularly focused in the OPL and RPE, although higher levels of expression in the GCL was observed in NHP compared to humans (Fig. 2C, D, Fig. S2E-H).

**Figure 2.**
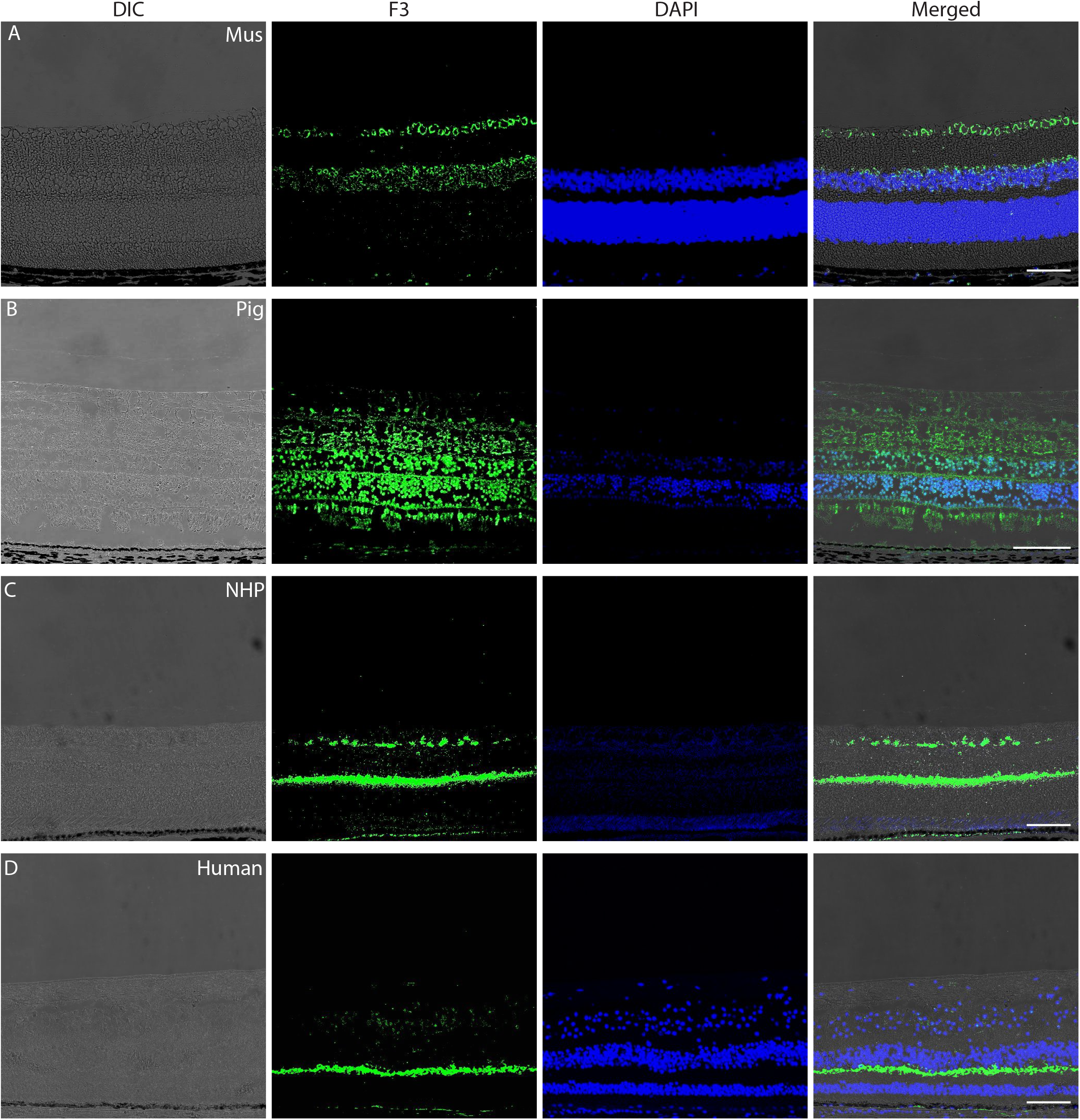
F3 distribution and localization in the retina across species. Representative images (n = 3-5) of cross-sections showing differential interface contrast (DIC), F3 expression (green), and nuclear stain (DAPI, blue) for (**A**) mus, (**B**) pig, (**C**) NHP, and healthy (**D**) human eyes. Scale bar = 100 μm.

Since F3 is also expressed more broadly throughout the eye than just the retina and RPE^26^, we also tested F3 staining in the cornea (Fig. S3A-D) and ciliary body/iris (Fig. S4A-D) in mice, pigs, NPHs and humans. Strong F3 staining was observed in the corneal epithelium and stroma of mice along with relatively weaker staining of the corneal endothelium (Fig. S3A). Similar degrees of epithelium staining were detected in human corneal epithelium (Fig. S3D). Pig and NHP demonstrated negligible F3 staining in the epithelium and stroma (Fig. S3B, C), however, it is important to emphasize here that these samples were more challenging to preserve during the fixation process. Very strong F3 expression was evident in the ciliary body and iris of mice (Fig. S4A), consistent with previous RNAScope analyses^15^. Similar, but weaker staining was observed in human ciliary body/iris (Fig. S4D), and very low, if any, F3 protein in pig and NHP ciliary body/iris tissue (Fig. S4B, C). Importantly, blocking peptide eliminated F3 staining in all ocular tissues tested (Fig. S5A-D), which, when combined with validation of the antibody in F3 KO mice (Fig. S1), confirm the specificity of our staining for F3 across species.

### F3 localization varies within the human retina and is associated with RPE deposits in AMD tissue

Until this point, we have used bulk RNA from specific ocular tissue to evaluate F3 expression, or relied on retinal F3 staining patterns within a single location, typically in the central retina adjacent to the optic nerve. However, retinal anatomy and physiology varies greatly depending on location. Thus, we decided to evaluate F3 staining throughout the retina, ora serrata to ora serrata (Fig.3A). In doing so, we detected surprising differences in F3 localization in human tissue. Peripheral retinal F3 staining (close to the ora serrata) was most intense in the RPE, followed by lower degrees of staining in the INL and ONL (Fig. 3B). As we moved inward towards the central retina (~3 mm from the optic nerve head), F3 staining began to shift from primarily RPE staining to predominantly OPL staining, with faint RPE and GCL/nerve fiber layer staining (Fig. 3C). Finally, within the foveal pit of the macula, F3 staining was predominantly in the OPL with faint staining in the INL and surprisingly devoid of positive F3 staining in the RPE (Fig. 3D). No such differences in F3 localization in the far-periphery, mid-periphery, or central retina were observed in mice or pigs (Fig. S6). We speculate that these differences might be present in the NHP retina, however, due to limited availability of ideal sections (entire retina with macula) we were unable to demonstrate this. Nevertheless, our results delineate another key difference observed in expression profile of F3 between mice and humans which is important for understanding basic F3 biology and eventually pathophysiology.

**Figure 3.**
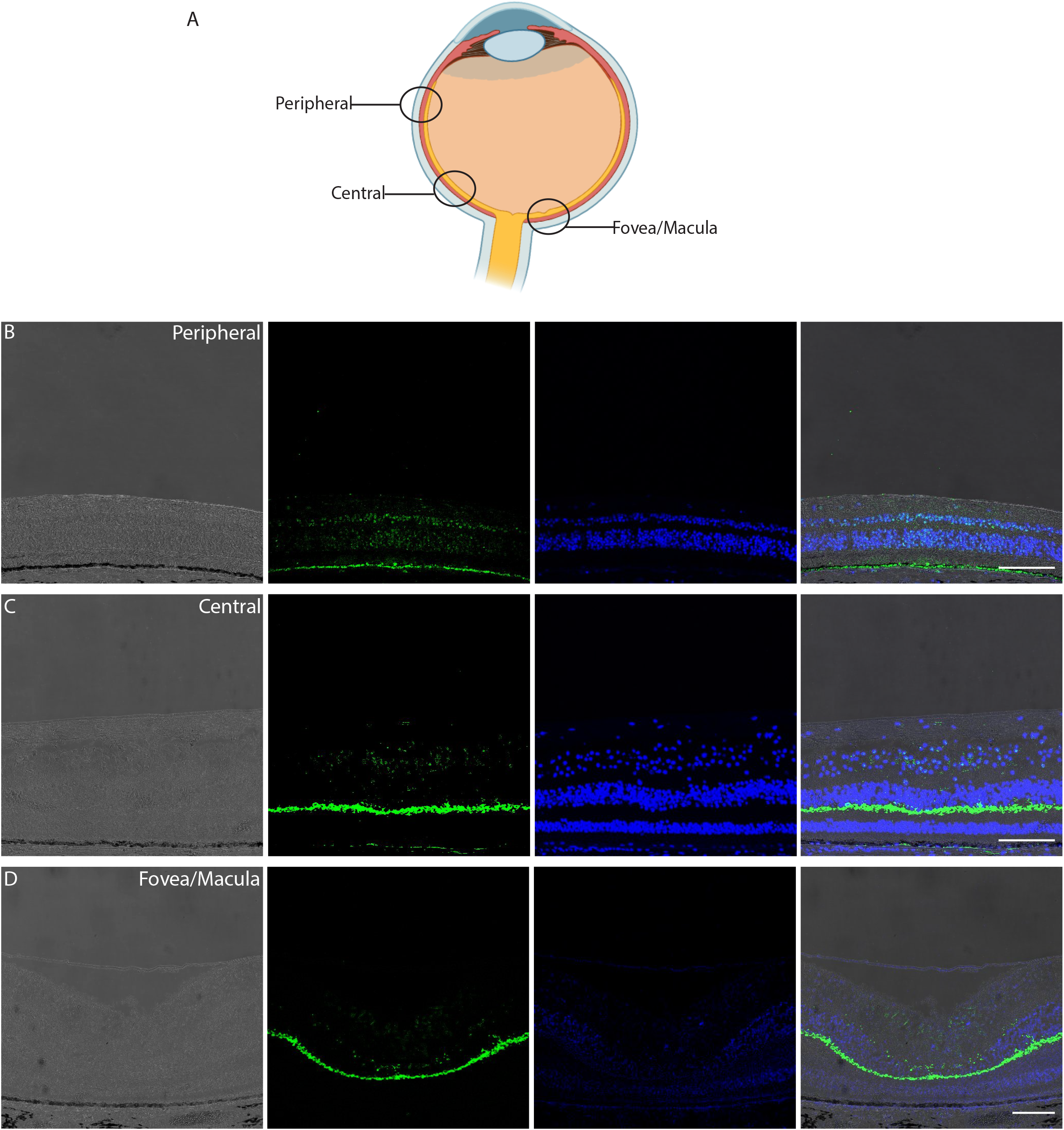
F3 localization varies within a human retina. (**A**) Schematic of the region of interest imaged in the human retina. Illustration from BioRender. Representative images (n = 3) of crosssections showing DIC, F3 expression (green), nuclear stain (DAPI, blue) for (**B**) peripheral (at ora serrata), (**C**) central (~3 mm from optic nerve head), and (**D**) fovea/macula region of a human retina. (scale bar = 100 μm).

Previous reports have suggested that F3 surrounds drusen in AMD patients^28^, ^29^. To confirm these results with our F3 KO-validated antibody, we stained ocular sections from two early AMD patients (Table S3, Fig. 4A, B). In fields of view that contained small drusen surrounded by sloughed off RPE cells (Fig. 4A, B), we confirmed positive F3 staining surrounding some, but not all small druse, consistent with previous reports^28, 29^. It is difficult to conclude whether the observed F3 staining is due to F3 expression inside the RPE, or if it is deposited, extracellular F3. Staining of F3 in the OPL within these sections suggests that that this field of view originates from the central retina.

**Figure 4.**
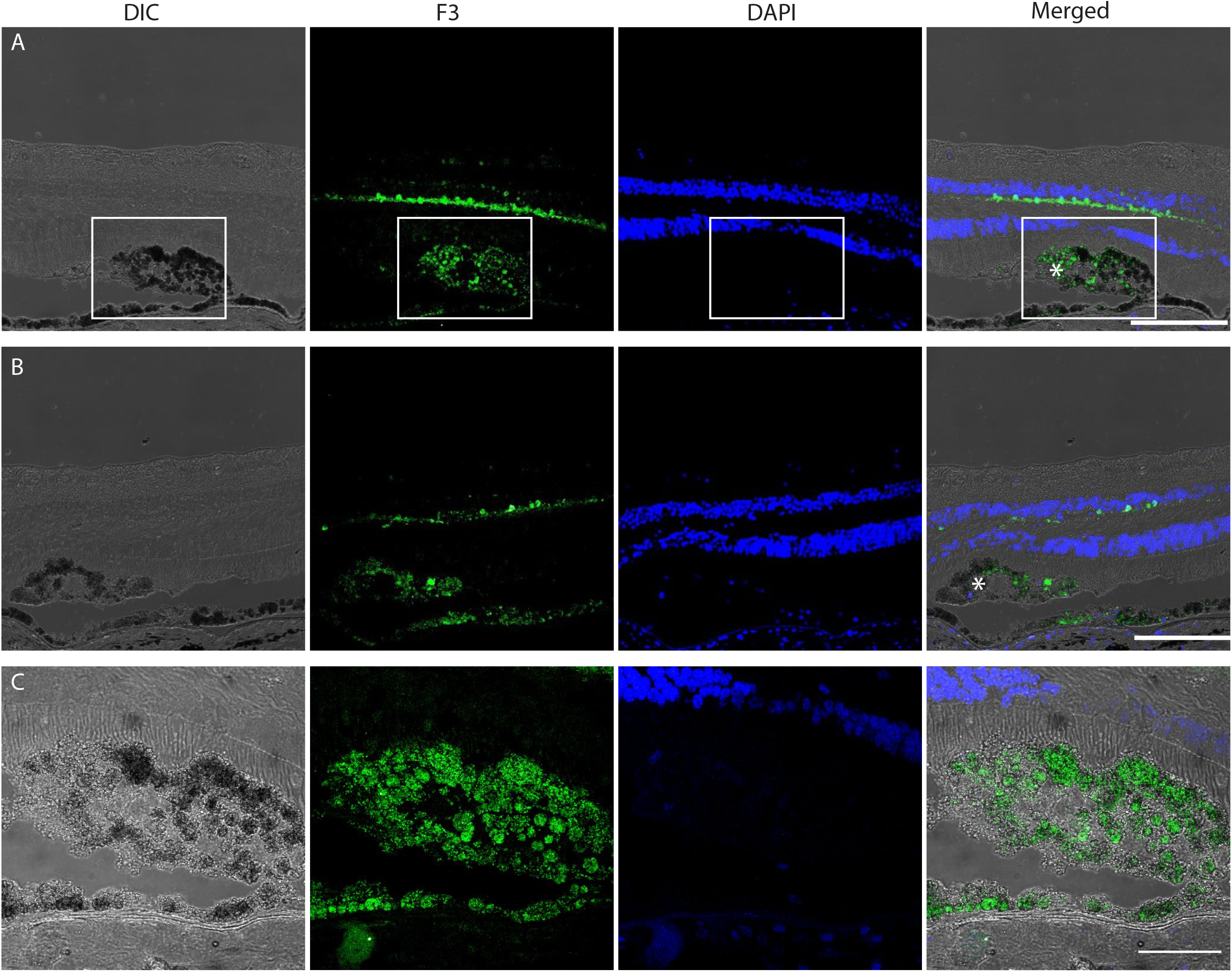
F3 is associated with RPE deposits in human eyes with early AMD. (**A-B**) Representative images of cross-sections showing DIC, F3 expression (green), nuclear stain DAPI (blue) for retinas of 2 early AMD patients focused on RPE pathology. Asterix (*) denotes F3 staining in small druse and surrounding F3 aggregation. (**C**) Zoomed-in image of druse and F3 accumulation from inset in (**A**) (scale bar = 100 μm).

### Aging alters F3 expression and ECM proteases involved in F3 turnover

The amount of F3 protein that we observe in any given tissue sample is a result of two primary factors, i) extent of F3 expression, and ii) F3 protein turnover. Recent studies have suggested that F3 expression increases with age^30^ and that F3 protein stability (in the form of amyloid) may also be increased with age^31^. In fact, F3 was identified as one of the top 5 genes overexpressed during aging across all analyzed tissues^30^. To determine whether F3 expression and protein levels also increases with age in the retina and RPE, we performed gene expression analysis in mice at 2-18 months of age (Fig. 5A) along with corresponding F3 immunostaining at similar ages of 4-19 months (Fig. 5B-D). As a control tissue for the expression experiments, we included RNA isolated from mouse liver. Surprisingly, we found that over the course of advance aging, F3 expression did not change in any tissue between 2 mo (young) and 6 mo (adult) of age in mice (Fig. 5A), but significantly increased between 6 mo and 18 mo (geriatric) mice in the NR (average of 10 fold) and liver (average of 25 fold) (p < 0.0001, Fig. 5A). F3 also increased in the RPE, but not significantly (Fig. 5A). Immunostaining for F3 confirmed the NR observations of increased protein levels in geriatric mice compared to young or adult mice (Fig. 5B-D).

**Figure 5.**
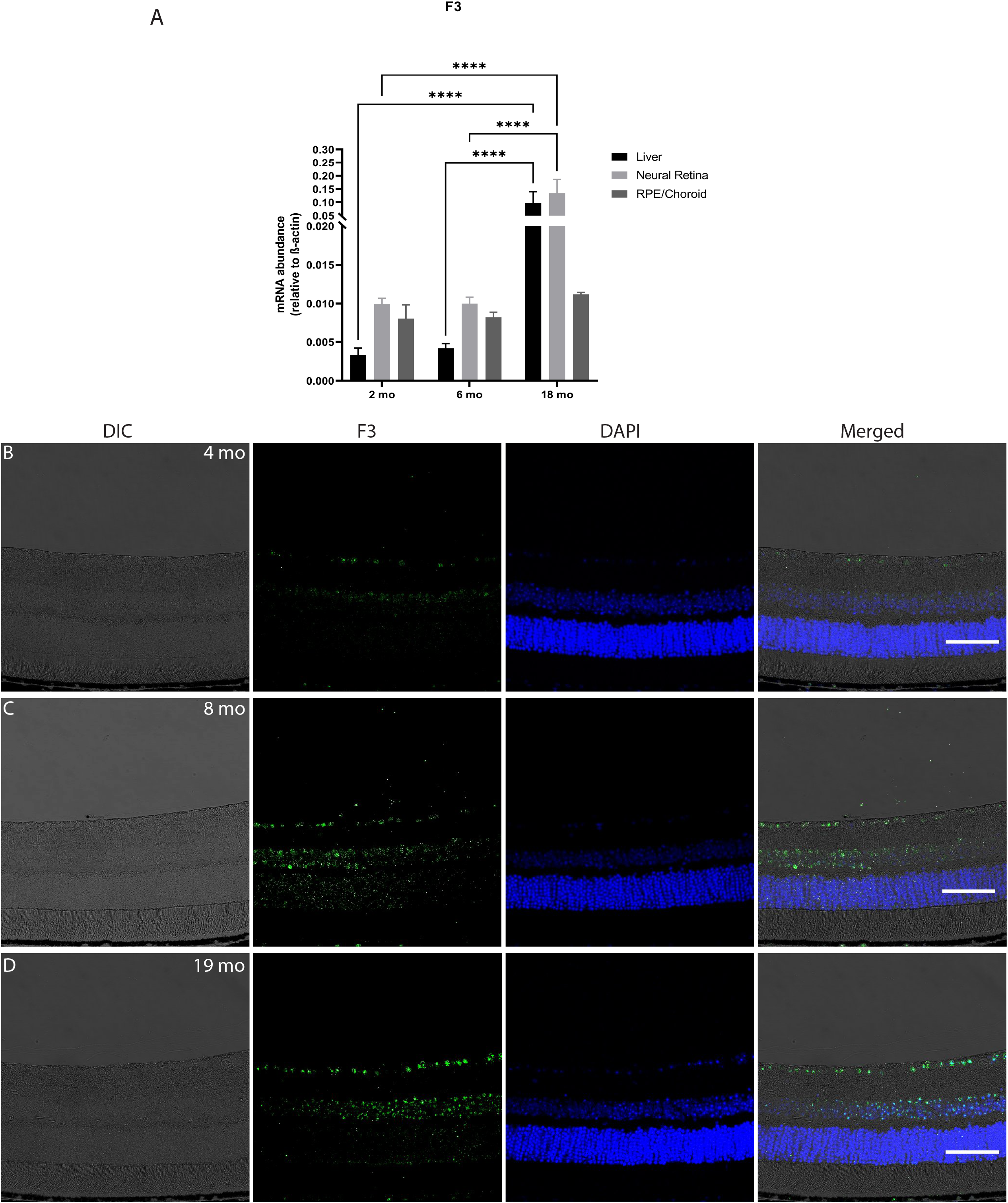
Age-related changes in F3 expression in mice. (**A**) Graph representing expression levels of F3 in the liver, NR and RPE of mice at 2 mo, 6 mo, and 18 mo relative to β-actin (n = 4, ****p<0.0001). Analyses performed by 2-way ANOVA and represented as mean ± SD. (**B**) Representative images (n = 4) of ocular cross-sections showing DIC, F3 expression (green), nuclear stain (DAPI, blue) of mice at 4 mo, 8 mo, and 19 mo. (scale bar = 100 μm).

While little is truly known about the totality of factors that regulate F3 protein turnover, two enzymes present in the eye that have been demonstrated to promote the degradation of F3 are matrix metalloproteinase 2 (MMP2)^32^ and high-temperature requirement A serine peptidase 1 (HTRA1)^33^. We next asked if the buildup of F3 that we observe with advanced age may be due in part to declining levels of these ECM homeostasis enzymes. qPCR was performed on two major ocular MMPs, Mmp2^34^ and Mmp9, and their cognate inhibitor, tissue inhibitor of matrix metalloproteinase 3 (Timp3)^35, 36^. With increasing age, we found a corresponding significant increase in Timp3 expression (p <0.0001, Fig. 6A) which was paralleled by a decrease in Mmp2 and Mmp9 expression (p < 0.001, p < 0.0001, Fig. 6B, C). Additionally, Htra1, a member of the secreted serine protease family linked which is linked to F3 turnover^33^ and AMD pathology^37^, decreases significantly in the NR as the mice age (p < 0.0001, Fig. 6D). Our data strongly suggest that age causes increases in F3 expression while also downregulating enzymes responsible for F3 extracellular degradation, the culmination of which promotes increased F3 production and deposition in the retina with advanced age, possibly promoting retinal dysfunction and age-related pathologies.

**Figure 6.**
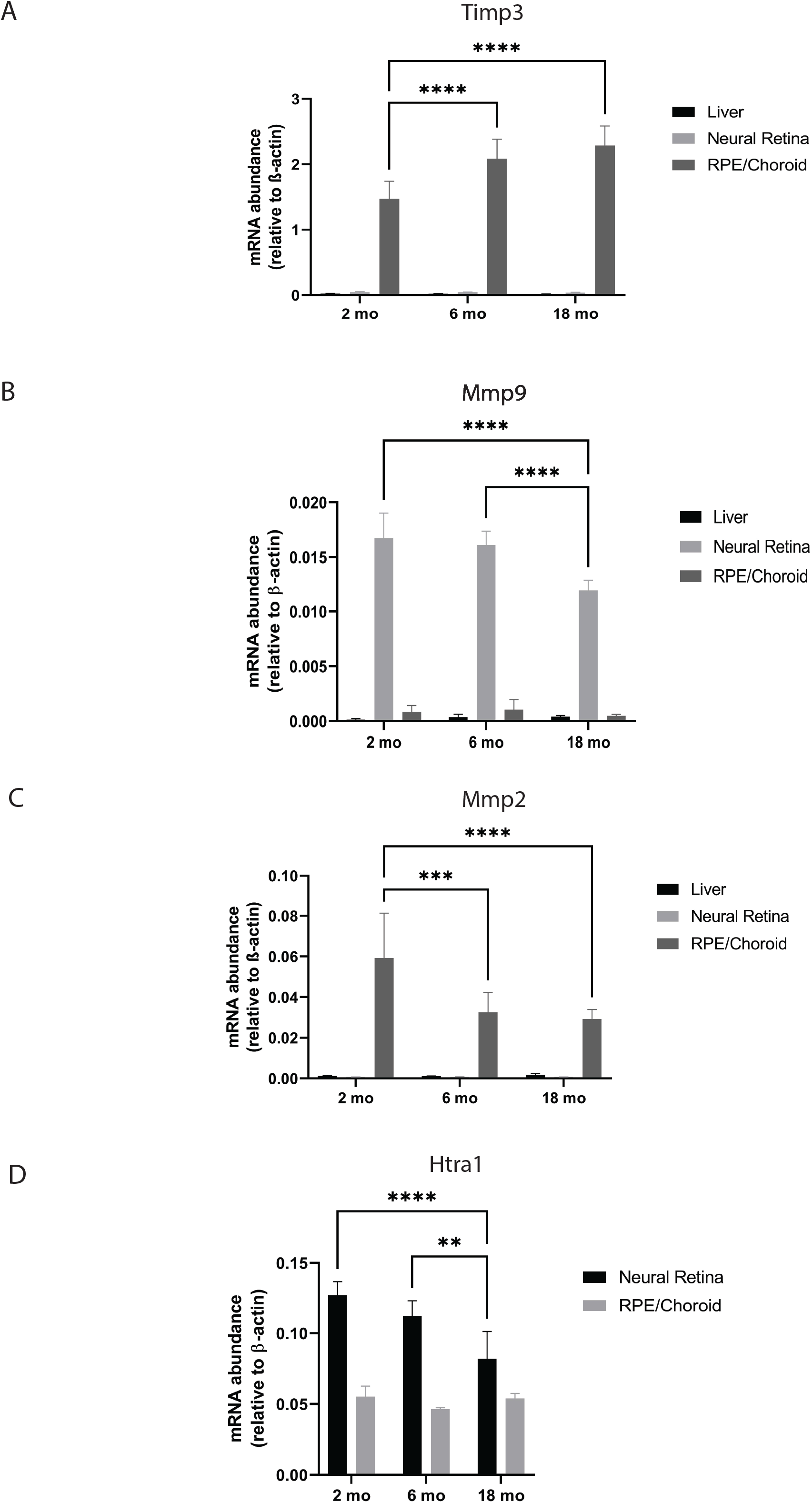
Age-related changes in the expression of ECM remodeling enzymes in mice. (**A**) Graph representing expression levels of Timp3 in the liver, NR and RPE of mice at 2 mo, 6 mo, and 18 mo relative to β-actin (n = 4, ****p<0.0001). (**B**) Graph representing expression levels of Mmmp9 in the liver, NR and RPE of mice at 2 mo, 6 mo, and 18 mo relative to β-actin (n = 4, ****p<0.0001). (**C**) Graph representing expression levels of Mmp2 in the liver, NR and RPE of mice at 2 mo, 6 mo, and 18 mo relative to β-actin (n = 4, ****p<0.0001, ***p=0.0003). (**D**) Graph representing expression levels of Htra1 in the liver, NR and RPE of mice at 2 mo, 6 mo, and 18 mo relative to β-actin (n = 4, ****p<0.0001, **p=0.003). All analyses performed by 2-way ANOVA and data are presented as mean ± SD.

## DISCUSSION

F3 is emerging as a functionally unique and physiologically important member of the fibulin family involved in several ocular pathologies in both the anterior and posterior chamber. Our results demonstrate the importance incorporating anatomical, age, and species related considerations when evaluating F3 expression and protein levels/localization in eye tissue. In addition to these factors, other studies have indicated that subject sex or sex hormones likely influence F3 expression^38, 39^ or BLamD formation^27^, although our studies were unable to confirm or refute those observations. The culmination of these data suggest that F3-based experiments must be appropriately thoroughly controlled and that observations made in lower order species, like mice, should be validated in separate, complementary model systems to enhance experimental rigor.

Our results provide a number of surprising and/or underappreciated observations regarding F3 behavior. First, the low level of RPE expression in mice relative to any other species tested was concerning. This is especially true since mice are currently the ‘go-to’ laboratory model for retinal diseases such as AMD wherein the primary pathology (drusen) is found between the RPE basal lamina and the inner collagenous layer of Bruch’s membrane. Indeed, R345W F3 knockin mice develop extensive and continuous BLamDs, but this phenotype takes 12-18 mo for robust development^19, 20^, and is typically concentrated in select quadrants of the eye^27^. It is interesting to speculate that identifying transgenic approaches aimed at elevating F3 RPE expression to levels observed in humans could be an approach to generate a more robust (or earlier onset) AMD-like mouse model. For example, using RPE-specific promoters (BEST1^40^ or RPE65^41^) could drive higher F3 levels in the RPE which may more closely match humanized F3 expression patterns. In addition, generating knockin mice harboring glaucoma-associated mutations in F3^14–16^ might provide an excellent glaucoma model, especially due to the high levels of F3 expression in the trabecular meshwork ring and ciliary body/iris.

Secondly, in the healthy human eye, we observed drastic changes in F3 protein localization in the retina/RPE depending on anatomical region – essentially favoring RPE accumulation in the peripheral retina, and OPL accumulation as one approaches the central retina and macula/fovea. One aspect that is surprising about these findings is the lack of apparent F3 RPE localization in areas that are susceptible to drusen formation (like the macula)^42^, especially in the case of DHRD/ML^18^. While the reasons for these distinct F3 localization changes is not clear, it is possible that there are F3 expression differences in retinal regions, or alterations in proteins that turnover F3 (e.g., MMP2 and/or HTRA1) in these particular regions that change with age and/or disease. Indeed, even in ocular cells that were once thought to be rather uniform, such as the RPE, more evidence suggests that there are physical and genetic differences that distinguish cell populations in the periphery vs. central retina vs. the macula^43^, and these results likely extend to additional retinal cell types too.

The primary staining of F3 within the fovea was restricted to the OPL, a retinal cell layer typically packed with neuronal synapses originating from horizonal cells, bipolar cells, and photoreceptors. According to single cell RNAseq (scRNAseq) data, human F3 is primarily produced in fibroblasts > RPE > Müller glia^44^. Thus, it is likely that the F3 we observe in the OPL originated from Müller glia, which span the height of the retina from the inner limiting membrane to the external limiting membrane. Due to their expansive nature, alterations in F3 protein patterns within the retina could reflect Müller glia F3 expression, but differential deposition due to altered F3 secretion or turnover patterns. Indeed, it is quite possible that the strong F3 staining we observe in the OPL may correlate to Müller glia endfeet^45^, or cone pedicles/spherules. Nonetheless, while we believe that F3 likely acts proximal to the cells that it is produced in, since it is a secreted protein, we cannot rule out that it may be produced in one location, but accumulates in locales distant from its source. Third, we demonstrate that F3 expression and protein levels in the mouse retina significantly increase with advanced age. The mechanisms that regulate F3 expression and turnover within the eye are virtually unstudied. Thus, it is difficult to even speculate what transcriptional regulatory pathways are involved in this age-dependent observation. Regardless, concomitant with the increase in F3 expression levels, we also noted significant reductions in the expression of proteins that have been demonstrated to proteolyze F3 in biochemical assays (Mmp2 and Htra1). These two findings highlight the likelihood of increased F3 deposition and dysfunction with age. Moreover, when these phenomena occur in the presence of an F3 mutation that causes misfolding^17, 46–48^, retinal disease could be further accelerated as an individual ages. However, given the vast differences we have observed in F3-based behavior between species, it is important that these results be reproduced in another species, ideally from human donor tissue.

As more studies have implicated F3 as a key player in disease, it has become more imperative to investigate this protein from a multidimensional standpoint to better understand its pathobiological relevance. Our findings shed light on important differences in F3 expression and localization in the eyes of a range of relevant species, providing an important baseline from which to compare new F3-based observations. Similarly, these data lay the groundwork for identifying and distinguishing ‘normal’ vs ‘diseased’ F3 characteristics, which is crucial for developing therapeutic interventions targeting this perplexing, yet fascinating, protein.

## ACKNOWLEDGEMENTS

We thank Dr. Christine Curcio, Ph.D. and Dr. Dongfeng Cao, Ph.D. (University of Alabama at Birmingham) for providing the early AMD donor slides. JDH is supported by an endowment from the Roger and Dorothy Hirl Research Fund and R01 EY027785. We thank the ARVO EyeFind Research Grant GAA202107-0016 (to SD) for financial support to procure control human donor eyes. Additional support was provided by a National Eye Institute Visual Science Core Grant (P30 EY030413, to the UT Southwestern Department of Ophthalmology).

**Sup. Figure 1. Comparison of commercially available F3 antibodies for specificity.**

Representative images of cross-sections showing merged (DIC/F3(green)/DAPI(blue) images for F3 wild-type (WT) and knockout (KO) mus (15 mo) using antibodies against F3 from ProSci (A&E), GeneTex (B&F), Chemicon (C&G), and Santa Cruz (D&H). Arrows indicate non-specific staining. (scale bar = 100 μm).

**Sup. Figure 2. Side-by-side comparison of F3 staining with corresponding H&E histology.**

(A, C, E, G) Representative images (n = 3-5) of cross-sections showing DIC, F3 expression (green), nuclear stain (DAPI, blue) for mus, pig, NHP, and healthy human eyes with their corresponding H&E (B, D, F, H) histology (GCL-ganglion cell layer, IPL-inner plexiform layer, INL-inner nuclear layer, OPL-outer plexiform layer, PR-photoreceptor, RPE-retinal pigment epithelium). (scale bar = 100 μm).

**Sup. Figure 3. F3 distribution and localization in the cornea across species.**

Representative images (n = 3-5) of cross-sections showing DIC, F3 expression (green), nuclear stain (DAPI, blue) for (**A**) mus, (**B**) pig, (**C**) NHP, and (**D**) human corneas. EP-epithelium, EN-endothelium. Scale bar = 100 μm [mus], 500 μm [pig, NHP, human]).

**Sup. Figure 4. F3 distribution and localization in the ciliary body and iris across species.**

Representative images (n = 3-5) of cross-sections showing DIC, F3 expression (green), nuclear stain (DAPI, blue) for (**A**) mus, (**B**) pig, (**C**) NHP, and (**D**) human ciliary body (CB) and iris (IR). (scale bar = 100 μm [mus] 500 μm [pig, NHP, human]).

**Sup. Figure 5. Absence of positive F3 staining in the presence of F3 blocking peptide demonstrates specificity of F3 antibody across tissues and species.**

Representative images (n = 3-5) of cross-sections showing DIC, F3 expression (green), nuclear stain (DAPI, blue) for (**A**) mus, (**B**) pig, (**C**) NHP, and (**D**) human-cornea, ciliary body, iris, and retina. Scale bar [cornea, ciliary body, and iris]= 100 μm, [mus] 500 μm, [pig, NHP, human]. Scale bar [retina]= 100 μm.

**Sup. Figure 6. Uniform F3 staining observed in different regions of mouse and pig retina.**

Representative images (n = 3-4) of cross-sections showing merged [DIC/F3 (green)/DAPI (blue)] for mus, pig retina. Scale bar = 100 μm.

