## Supplementary figures and images for "Important anatomical, age-related, and species considerations regarding ocular fibulin-3 (EFEMP1) analysis"

### Fig. S1

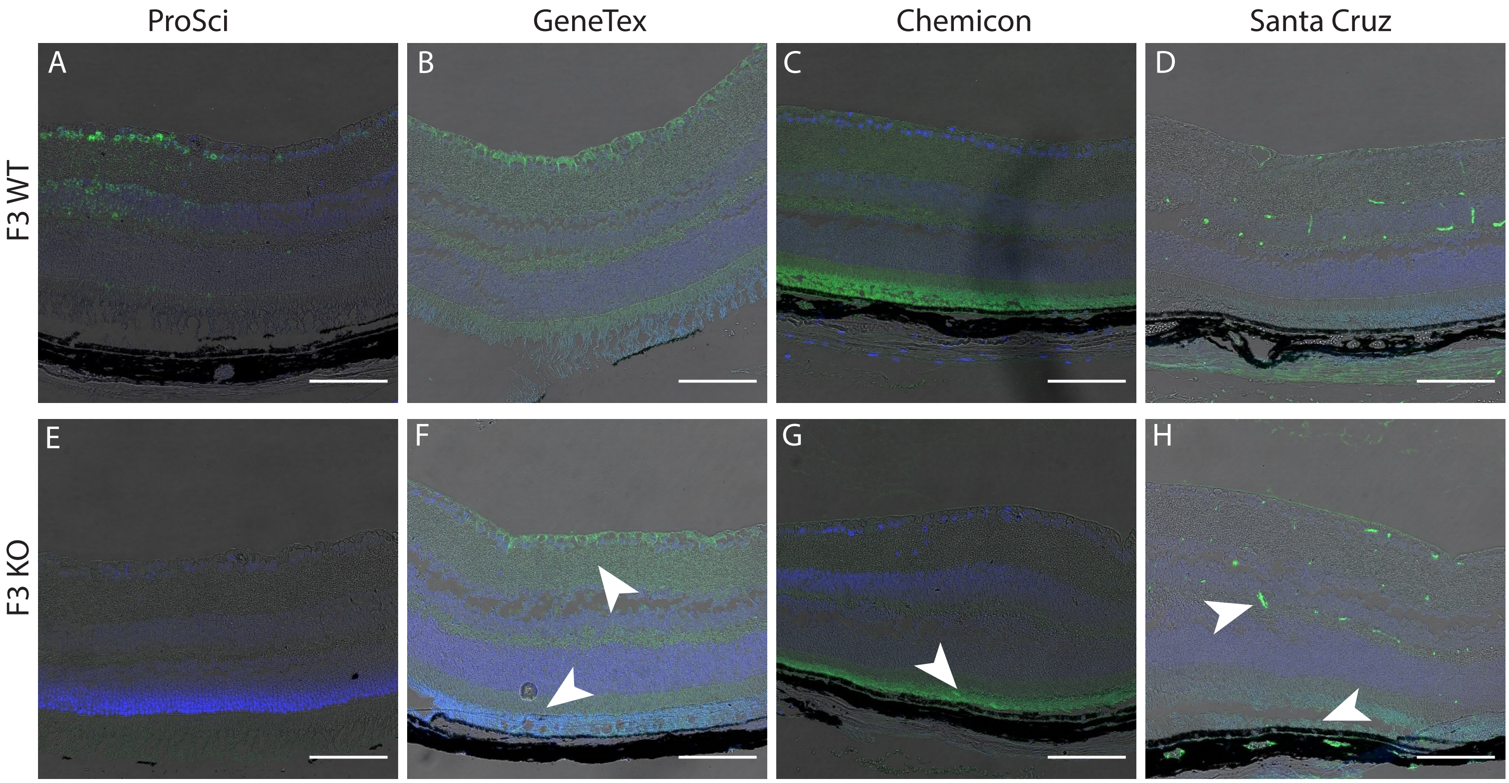


Sup.Fig.1

### Fig. S2

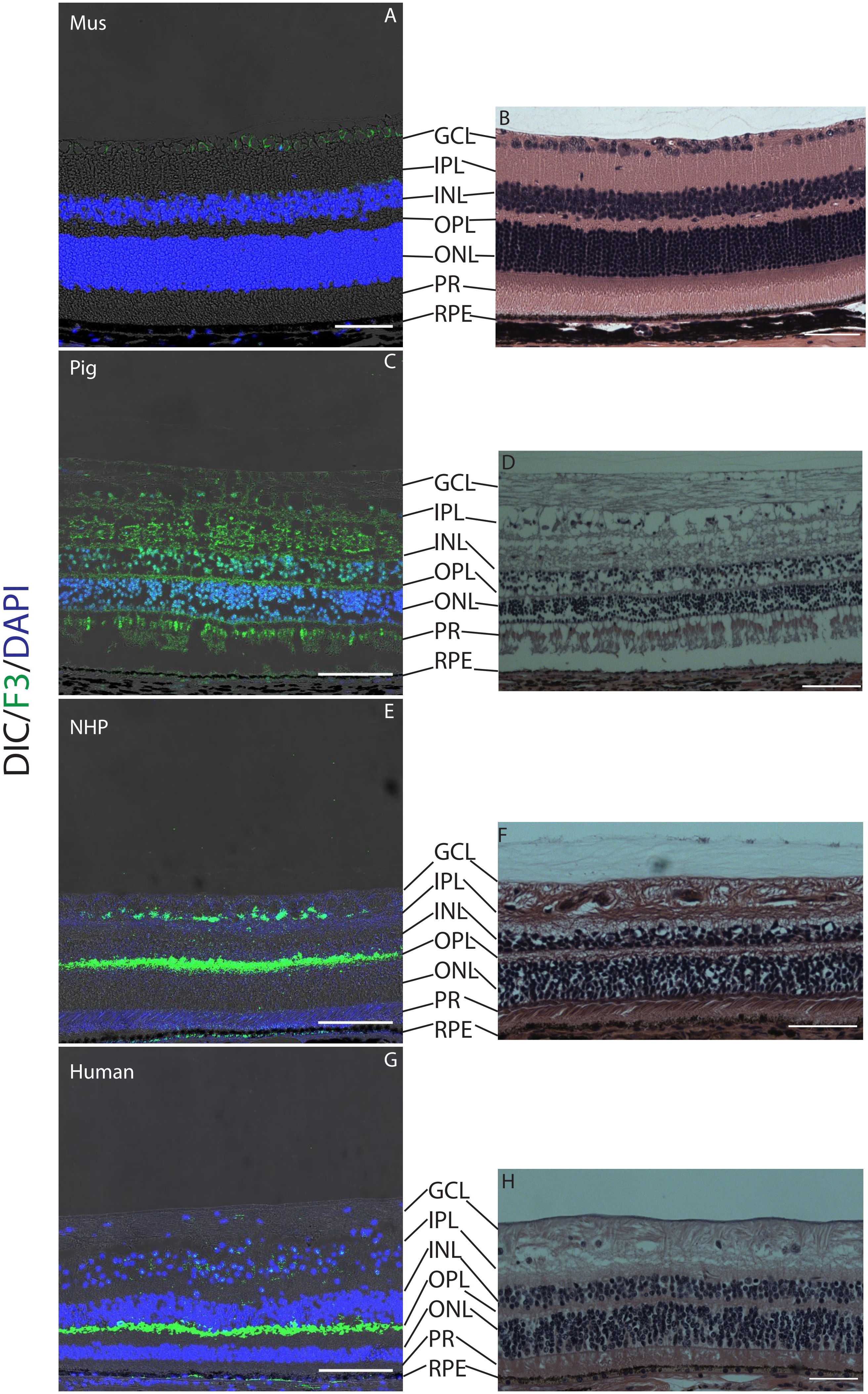

### Fig. S3

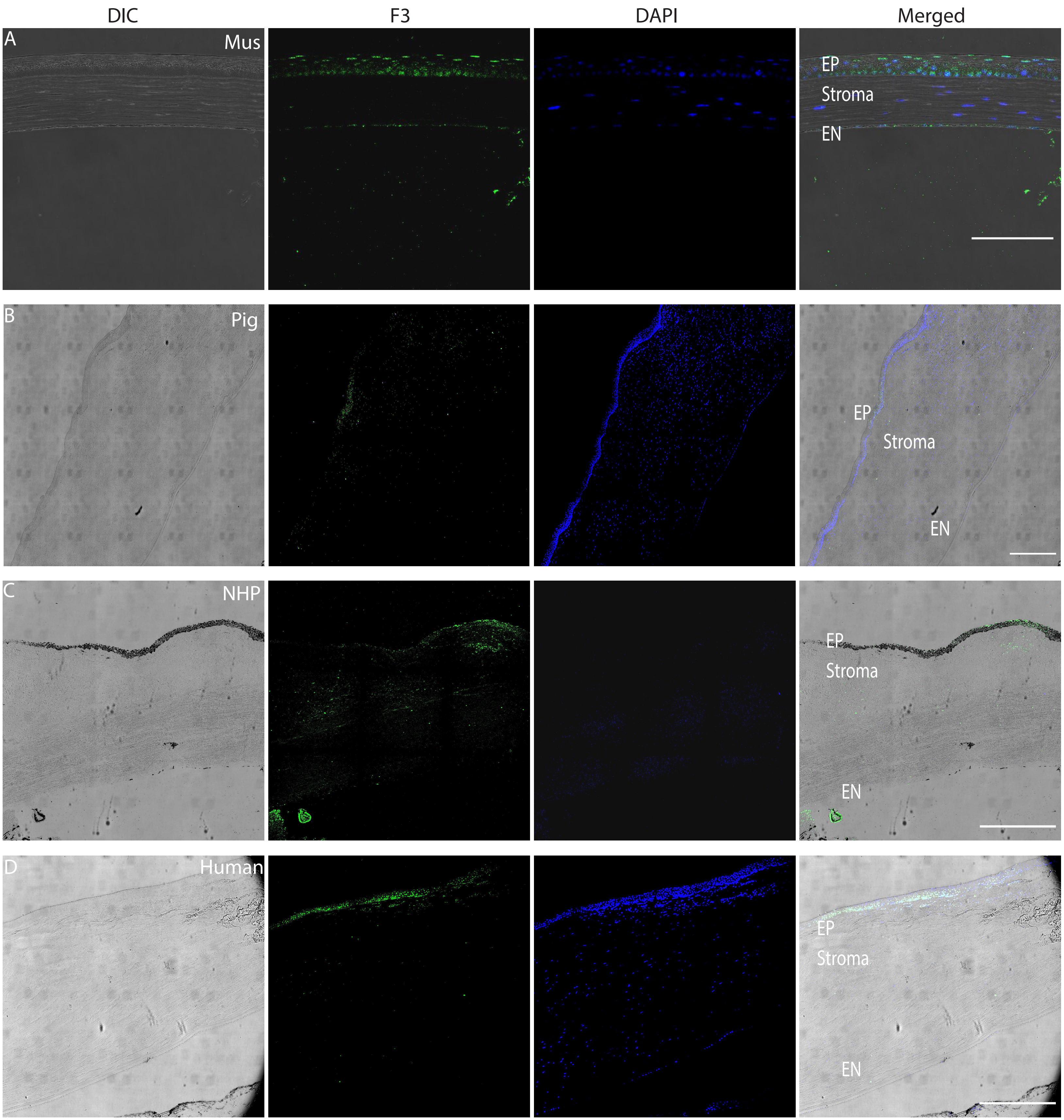

### Fig. S4

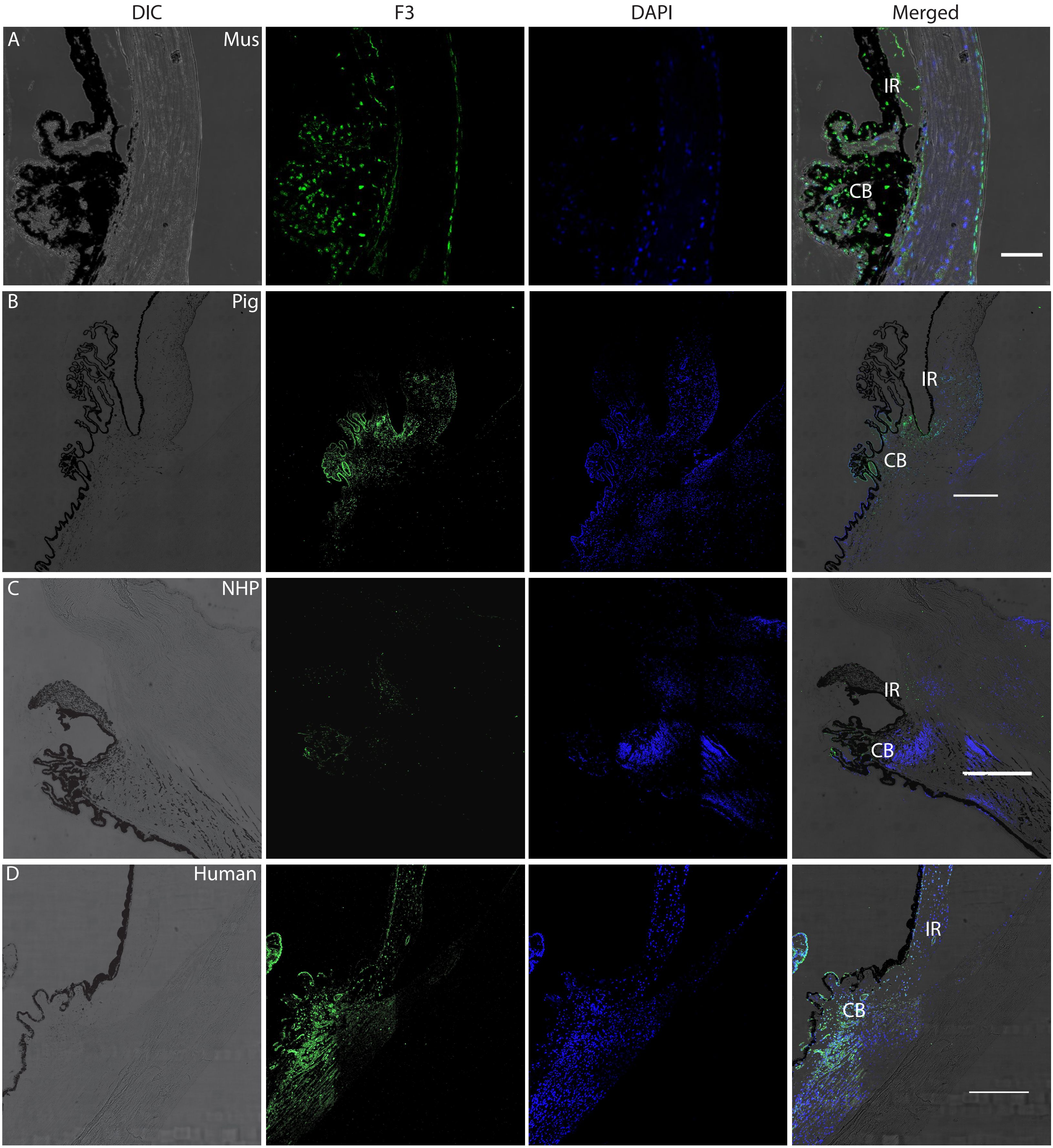


Sup.Fig.4

### Fig. S5

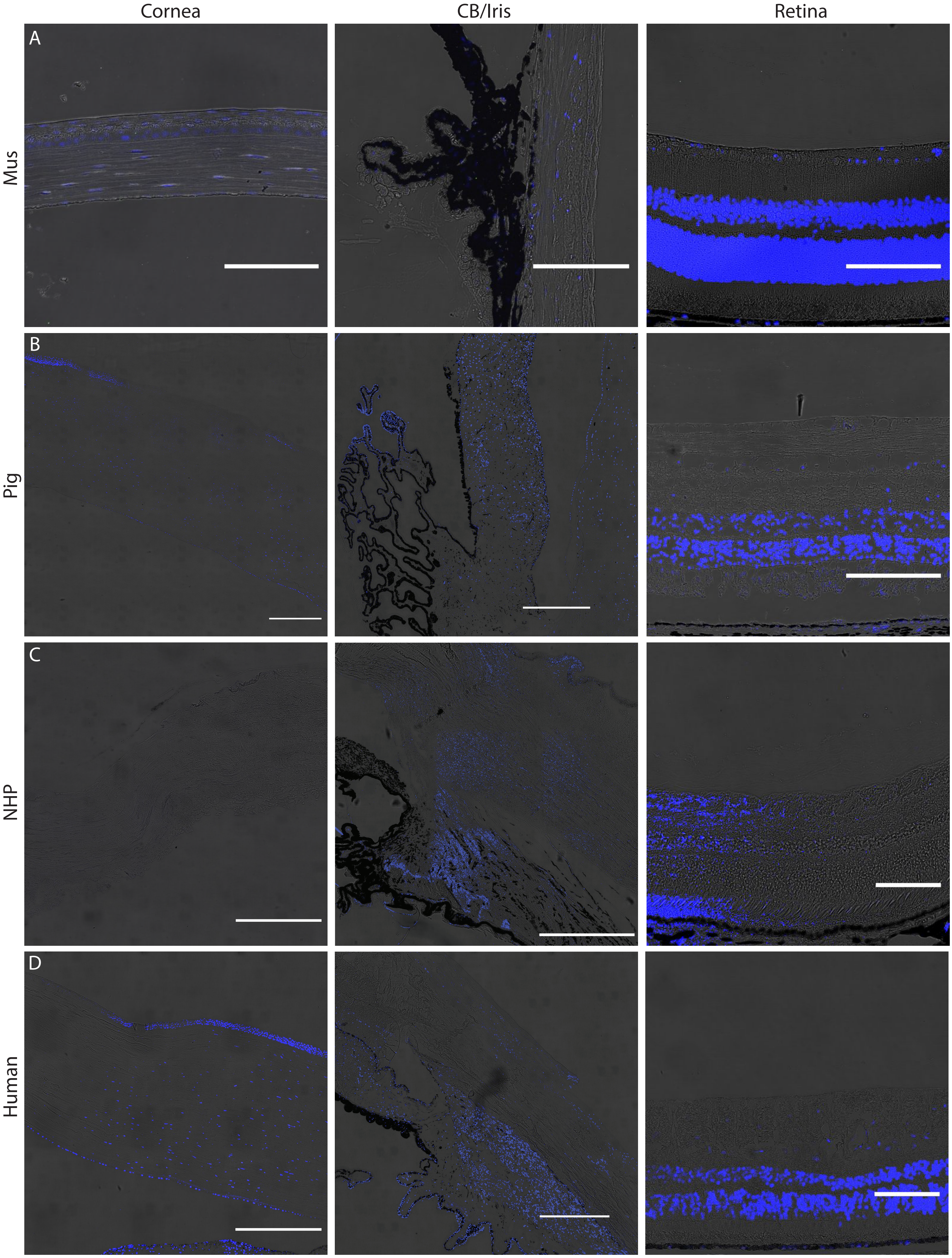


Sup.Fig.5

### Fig. S6

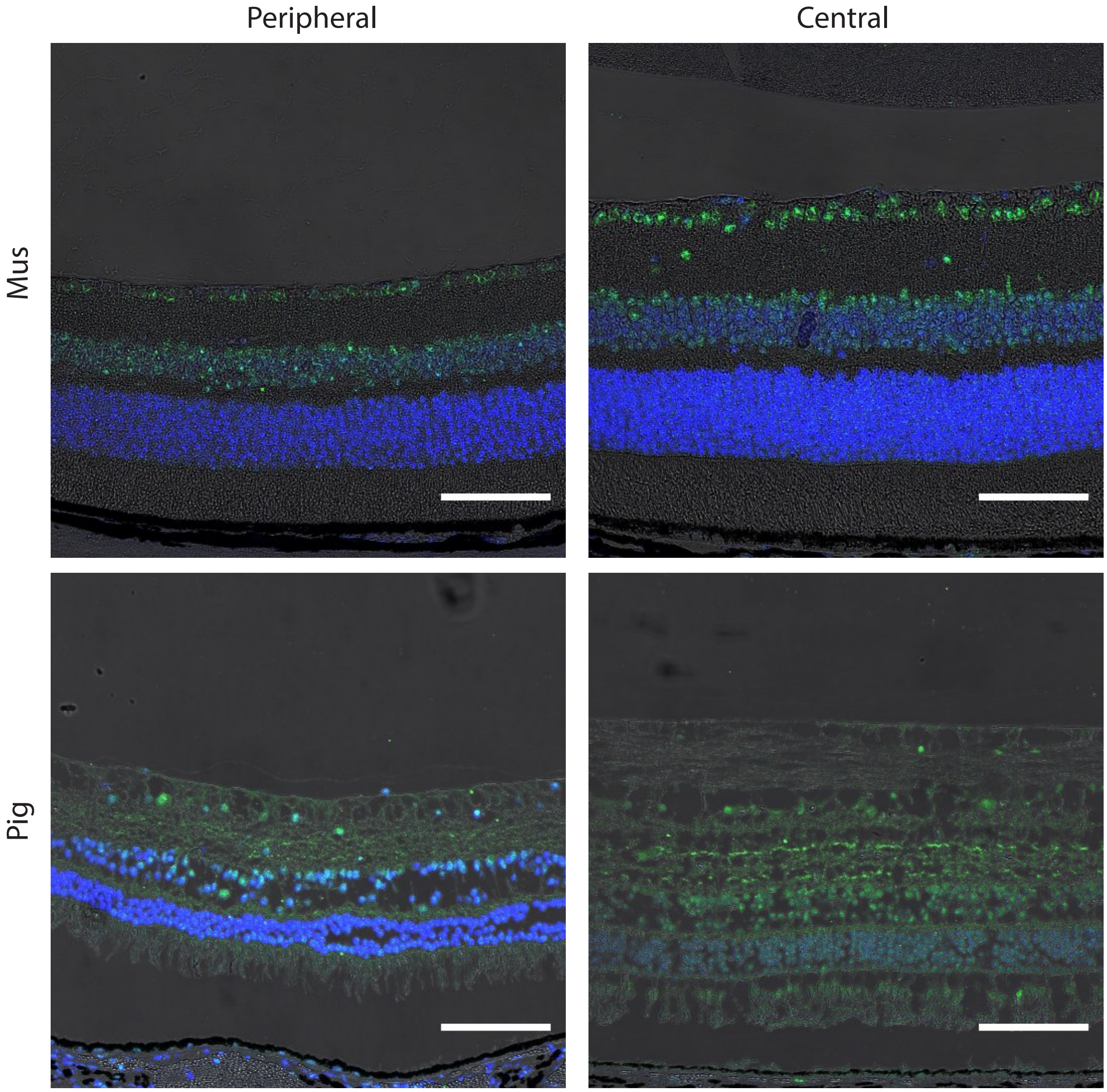


Sup.Fig.6
