## Supplementary material for "Important anatomical, age-related, and species considerations regarding ocular fibulin-3 (EFEMP1) analysis": Table S1-S3

**NHP**

| Sex | Species Code | Common Name | Scientific Name | Age (yrs) |
|---|---|---|---|---|
| F | PCX | Olive/Yellow Baboon | P.h. anubis/ P.h. cynocephalus | 1 |
| F | PCA | Olive Baboon | Papio hamadryas anubis | 12 |
| F | PCA | Olive Baboon | Papio hamadryas anubis | 10 |
| M | PCA | Olive Baboon | Papio hamadryas anubis | 2 |

**Supplemental table 1**. NHP donor information (OD used for RNA isolation, OS used for immunohistochemistry).

| Human | (Normal) | | |
|---|---|---|---|
| Sex | Race | Age (yrs) | Cause of death |
| F | Caucasian | 66 | unspecified |
| M | Caucasian | 85 | Cardiac arrest |
| M | Caucasian | 73 | Unspecified |
| F | Caucasian | 68 | Unspecified |
| M | Caucasian | 61 | Complications from heart bypass |

**Supplemental table 2**. Normal human donor information (OD used for RNA isolation, OS used for immunohistochemistry).

Human (AMD)

| Sex | Race | Age (yrs) | Cause of death | AMD category |
|---|---|---|---|---|
| F | Caucasian | 89 | Cholelithiasis | Early |
| M | Caucasian | 92 | Fall | Early |

**Supplemental table 3**. AMD donor information (OS used for immunohistochemistry).
